# Assessing the effects of female protections on size structure and spawning potential in two clawed lobster fisheries subject to varying exploitation levels

**DOI:** 10.1101/2024.08.05.606679

**Authors:** Kaitlyn Theberge, Tonje K. Sørdalen, Tracy L. Pugh, Holly K. Kindsvater

## Abstract

Current fishery management practices in both the U.S. Gulf of Maine and Norwegian clawed lobster fisheries primarily focus on conserving mature females to maximize egg production. While abundance of adult American lobsters (*Homarus americanus*) in the Gulf of Maine remains high, declines appear to be on the horizon. Similarly, the European lobster (*Homarus gammarus*), is facing its lowest recorded population size in southern Norway. Understanding how management strategies and fishing practices impact lobster size structure and spawning potential could inform management to improve resiliency to climate-induced changes. In the Gulf of Maine fishery, egg-bearing (ovigerous) female lobsters are not only protected from harvest, but also v-notched which offers additional protection up to several years. Norway, however, protects egg-bearing females without v-notching. Comparing these fisheries allows us to test the effect of the different management practices and how they interact with key vital rates, including growth and natural mortality rates. We used deterministic size- and age-structured models and empirically estimated growth and molt functions to simulate relative changes in abundance, size structure, egg production, and sex ratios in response to <u>these</u> two female protection strategies. Our findings suggest that in all scenarios, controlling total fishing effort to low or moderate levels - relative to the *F* > 1 that has been estimated for American lobster - is most important for the effectiveness of size-based restrictions on harvest of larger individuals. Both forms of female protection enhance overall egg production in both species across levels of fishing intensity, but also result in a skewed sex ratio in favor of females and a more pronounced size disparity between female and male lobsters. Moreover, our results suggest that American and European lobster populations exhibit differential responses to the management strategies, likely due to variations in estimates of natural mortality rates and growth rates. Our results highlight the sensitivity of management effectiveness to assumptions regarding the underlying biology, but also provide a clear message that current intense fishing practices have likely depleted the ability of both species to compensate for fishing mortality in the long term.

## 1. INTRODUCTION

Practices to improve the sustainable management of marine fisheries include regulations for the size and sex of legal catch. While the intention of these regulations is often to protect older mature females, they can also have unintended consequences for various aspects of population dynamics, including size structure, sex ratio, and mating behavior (Hines *et al*. 2003; Rowe and Hutchings 2003; Sato *et al*. 2005; Baker *et al*. 2022). These downstream effects can impact recruitment, ultimately either supporting or undermining population productivity. Consequently, discerning how populations respond demographically to selective harvesting remains essential for forecasting sustainability and designing optimal management strategies (Rochet and Marty 2016). Crustacean fisheries worldwide use a diverse range of management regulations, in part due to the distinctive characteristics of lobsters and crabs, which allow non-invasive sex identification and because females carry their developing eggs externally. For instance, lobster fisheries often implement bans on the harvesting of females within the harvestable size range if the female is carrying eggs, with the goal of enhancing overall egg production as a countermeasure against overfishing.

American lobsters (*Homarus americanus*) are widely distributed in the northwestern Atlantic Ocean from Newfoundland, Canada to North Carolina, USA; however most are found north of Rhode Island, USA (Caputi *et al*. 2013). European lobsters (*Homarus gammarus*) are found in the northeastern Atlantic from the Norwegian Arctic Circle to as far south as Morocco and parts of the Mediterranean (Wahle *et al*. 2013). American and European lobsters are close relatives, and share many biological similarities, but the fisheries for each species are managed differently to account for fishery scale and local cultural values. Contrasting the American lobster fishery in the U.S. Gulf of Maine to the European lobster fishery in southern Norway provides an opportunity to more closely examine management strategies’ effects on demography. This includes exploring how life history traits like size at maturity or natural mortality influence the consequences of management on sex-specific size structure, sex ratio, and annual egg production.

In the U. S. Gulf of Maine American lobster fishery, the harvesting of egg-bearing females is prohibited. Fishers are also required to make a V-shaped incision in the egg-bearing females’ tail, a practice known as “v-notching,” before releasing them back into the water (Code of Federal Regulations 2020); this protects the female from future harvest even while not carrying eggs. In the nearshore Gulf of Maine, where the majority of US landings originate, v-notching affords protection for up to four years (see Mazur *et al*. 2019). V-notching is not a common practice in Europe, and it is mainly found in Irish fisheries (Department of the Marine, Dublin 1994). In the Norwegian fishery for European lobster, harvesting of egg-bearing females is also prohibited, although it does not involve a v-notching requirement (Kleiven *et al*. 2011). Since the Norwegian fishery is open only for a two-month-long season (October and November), we assume this regulation provides only a one-year protection period for the egg-bearing females.

Protection of mature females coupled with elevated fishing pressure on males could affect adult sex ratios and sex-specific size structure. Past work on clawed lobsters suggests that differential fishing selectivity for males and females may lead to suboptimal mating conditions including skewed sex ratios (Daniel *et al*. 1989; Pugh *et al*. 2013; Jury *et al*. 2019), reduced mating success (Pugh 2014; Goldstein *et al*. 2014), and sperm limitation (Gosselin *et al*. 2005). It may also affect the prevalence of sexually-selected traits like claw size in males (Sørdalen *et al*. 2020). American lobster sex ratios (female:male) in coastal Maine have reportedly ranged from 1:1 in survey work to 1.4:1 in commercial traps in the 1970s and 1980s (Krouse 1973; Daniel *et al*. 1989). More recently, work in northern Massachusetts documented roughly equal sex ratios in smaller lobsters from commercial traps, but above harvestable sizes, the majority (60-90%) were female (Pugh *et al*. 2013). Differential protection of females, resulting in intense fishing pressure on males, has been suggested as at least partially responsible for female-skewed sex ratios in these studies as well as in other portions of the U.S. fishery (Jury *et al*. 2019).

The aim of this study is to identify how fishing intensity and types of female protections affect sex ratio, size structure, size disparity between females and males, and egg production for both American and European lobsters. We use demographic models of population dynamics to incorporate life history traits to predict population-level trends in size structure and productivity across a range of fishing pressures and management scenarios. Comparing both American and European lobsters in this study provides an opportunity to examine how two forms of restrictions on female harvest (a ban on harvest of egg-bearing females and v-notching, corresponding to one or four years of protection), interact with species life histories to impact population demographics.

## 2. METHODS

### 2.1 Model Overview

For both American and European lobster, we constructed a deterministic sex-specific population dynamics model using the open-source statistical software R version 4.0.2 (R Core Team 2020). This model was based on the methods for a deterministic model of age-structured population dynamics with annual time steps developed in Mangel (2006) and Kindsvater *et al*. (2020). We extended the model to incorporate the unique growth patterns of clawed lobsters in order to estimate average size at age, allowing us to address the size-selectivity of each fishery (Table 1), including variations on female lobster protections with slot limits (minimum and maximum sizes) on harvest. This approach made it possible to compare relative trends and changes in size structure, sex ratio, and total egg production for different scenarios of female protection in the American and European lobster populations. These attributes were chosen because they are quantifiable traits that are important for reproductive success. Furthermore, our approach provides an opportunity to examine relative trends in egg production and size structure for a range of fishing intensities, without the need for indices of abundance or landings data. The model was run for three scenarios of differing female protection (none, 1-year, 4-year), each at varying levels of fishing intensity (*F*) (see Figure 1). We included the southern Norwegian practice of banning harvest of ovigerous females which results in protection from harvest for one year (1-year protection scenario in the model), as well as the Gulf of Maine fishery requirement of v-notching egg-bearing females, assuming it protects them for four years, even if they are not carrying eggs (4-year protection scenario in the model). By comparing management strategies across a range of fishing pressures, and accounting for the unique patterns of growth and natural mortality that have been estimated for each species, we can predict the relative impacts of female protection on the size structure of the adult population, as well as resulting size disparities between the sexes for both clawed lobster fisheries.

**Figure 1.**
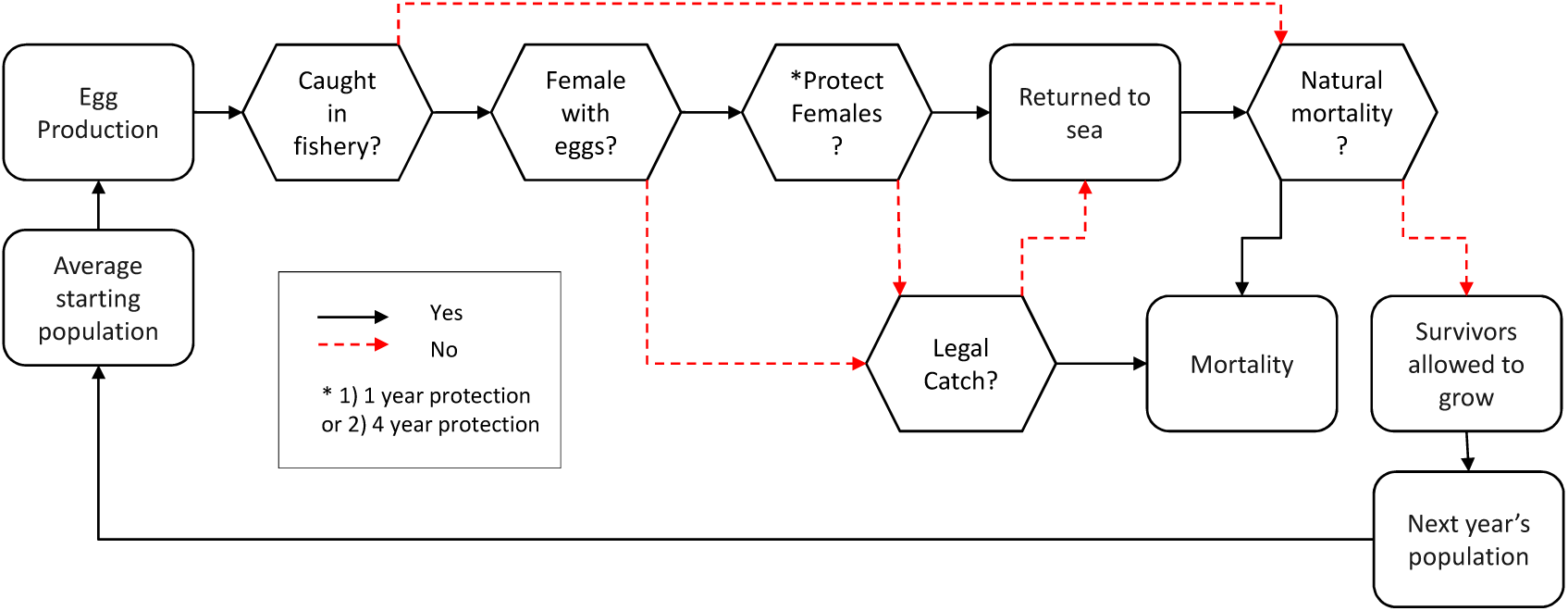
Summary of the order of events that the model simulates in each time step. Items with an asterisk are variable depending on the scenario.

**Table 1.** Harvest regulations for American (ASMFC 2020) and European lobster fisheries (Kleiven et al. 2011; Sørdalen et al. 2018).

| Regulation | Gulf of Maine American lobster (*H. americanus*) | Southern Norway European lobster (*H. gammarus*) |
| --- | --- | --- |
| Minimum harvestable size | 83 mm CL | **Females:** 86.9 mm CL<br>**Males:** 88.6 mm CL<br>(Converted from 250 mm total length for each sex) |
| Maximum harvestable size | 127 mm CL | **Females:** 114.76 mm CL<br>**Males:** 117.09 mm CL<br>(Converted from 320 mm total length) |
| Harvest season | January 1 – December 31 (all year) | October 1 – November 30 (Thorbjørnsen *et al.* 2018) |
| Trap limit | 800 traps per fisher, limited to 600 traps in Zone E (Alden 2023) | Commercial fishers may use up to 100 traps per day, whereas recreational fishers may use 10 traps per day |
| Egg-bearing female protection | V-notching, no harvest of egg-bearing females | No harvest of egg-bearing females |

### 2.2 Growth

Lobsters and other crustaceans have a discontinuous growth pattern where they undergo ecdysis, or molting. Lobsters molt more frequently when they are smaller and growing fast, and less frequently as they get older and larger (Phillips *et al*. 1980; Huntsberger *et al*. 2020). While recent advances have been made in aging crustaceans, the loss at each molt of the hard body parts that may track age makes it difficult to accurately assign age (Kilada *et al*. 2012; Wahle *et al*. 2013; Huntsberger *et al*. 2020; Fairfield *et al*. 2021). Thus lobster size-at-age models are based on data for molting increments (the difference between pre- and post-molt size) and the duration of time between molts (the intermolt period), which are typically collected from mark-recapture studies of wild individuals or laboratory growth experiments (Ennis 1978; Krouse 1981; Campbell 1983; Lawton *et al*. 1984; ASMFC 2020; Sørdalen *et al*. 2022).

Our estimates of size-at-age were based on previously compiled data for annual size-based molt frequency and molt increments gained from mark-recapture studies of both American (ASMFC 2006) and European lobster (Sørdalen *et al*. 2022). This allowed for a projection of the mean probabilistic relationship between size and age for each species. The model used estimates of mean annual growth to calculate egg production based on the mean size of spawning females. It also calculated mean change in size for males and females across scenarios. The growth equations for European lobster were parameterized using data from a tag-recapture survey from both protected areas and fishing areas in southern Norway (Sørdalen *et al*. 2022). Data on molt frequency for the largest individuals was unfortunately limited for both species.

Annual growth in terms of carapace length was the product of annual molting probability and molt increment. The annual probability of molting *P_molt_* and the molt increment *MI_g_* were calculated as a function of sex and pre-molt carapace length *CL*. Due to differences in the reporting practices for each fishery for each species (total length vs. carapace length), we converted total length to carapace length for European lobster, using the equations in Table 2. The frequency of molting and the increment of growth vary with individual size, temperature and between fished and unfished populations, which likely differ in density. Unfortunately, sufficient data to characterize these potential differences are not available.

**Table 2.** Species-specific relationships used to calculate average growth, maturation, and recruitment. Average growth per year is calculated by converting Total Length to Carapace Length (for European lobster), and then estimating the product of the probability of molting and the molt increment at each size.

| Parameter / Equation Name | Gulf of Maine American lobster (*H. americanus*) | Southern Norway European lobster (*H. gammarus*) |
| --- | --- | --- |
| Total Length (TL) to Carapace Length (CL) conversion | n/a | Female:<br>$CL = 0.3842 \times TL - 8.45$<br>Male:<br>$CL = 0.3987 \times TL - 10.42$<br>(Sørdalen *et al.* 2022) |
| Probability of molting ($P_{molt}$) | $p_{molt\ a} = \frac{1}{1 + e^{(a+b \times CL)}}$<br>$a$ = Female: -6.571<br>Male: -6.834<br>$b$ = Female: 0.05901<br>Male: 0.06046<br>(ASMFC 2009, 2020) | See Supplemental Table 1 |
| Molt Increment (MI) | $MI_a = \begin{cases} m_a + n_a \times CL & \text{for } CL < f \\ m_a + n_a f & \text{for } CL \geq f \end{cases}$<br>$m_a$ = Female: -3.90<br>Male: -4.51<br>$n_a$ = Female: 0.22<br>Male: 0.23<br>$f$ = Female: 76<br>Male: 86<br>(ASMFC 2009, 2020) | $MI_g = m_g + n_g \times CL$<br>$m_g$ = Female: 13.31382<br>Male: 9.2874<br>$n_g$ = Female: -0.05861<br>Male: 0.02223<br>(Sørdalen *et al.* 2022) |
| Maturation ogive parameters (Eqs 1 and 2) | $\alpha$ = -17.14056<br>$\beta$ = 0.19664<br>(ASMFC 2020) | $\alpha$ = -40.59<br>$\beta$ = 0.43<br>(Coleman *et al.* 2023) |
| $L_{mat}$ | 87 mm<br>(ASMFC 2020) | 92.5 mm<br>(Coleman *et al.* 2023) |
| Spawning frequency (SF) | $\begin{aligned} &\text{for } L_{mat} > CL > 120 \quad freq = 0.5 \\ &\text{for } CL > 120 \quad freq = 0.66 \end{aligned}$<br>(Waddy and Aiken 1986; Waddy *et al.* 1995; ASMFC 2020) | $freq = 0.50$<br>(Sørdalen 2019) |
| Fecundity at size (Y) | $Y = 1.09886(0.000919833[CL^{3.580220}])$<br>(Estrella and Cadrin 1995) | $Y = 468.17(CL - 33004)$<br>(Agnalt 2008) |
| Stock-Recruitment relationship parameters | $c = 1$<br>$d = 9E-10$ | $g = 100,000$ recruits per $10^8$ eggs<br>$h = 0.851$ eggs ($\times 10^{-8}$)<br>(Bannister and Addison 1986) |
| Instantaneous Natural Mortality $M$ | 0.15<br>(ASMFC 2020) | **Females:** *Small*: 0.24208<br>*Large*: 0.33281<br>**Males:** *Small*: 0.38074<br>*Large*: 0.63534<br>(Fernández-Chacón *et al.* 2021) |
| Size cutoff value for small and large natural mortality | n/a | **Females:** 88.16 mm CL<br>**Males:** 89.09 mm CL<br>(Converted from 250 mm TL)<br>(Fernández-Chacón *et al.* 2021) |

For American lobster, annual probability of molting was given by a logistic equation, and molt increment was a linear function until it reaches an inflection point, after which molt increments were constant (Table 2). Note that *f* represented an inflection point where the length at which 10% of individuals were mature and presumably beginning to allocate resources to reproduction instead of growth (ASMFC 2009, 2020). Annual probability of molting was then multiplied by the molt increment for each size class to calculate the mean annual growth for a given sex and pre-molt size. Molt increment for European lobster females and males were linear functions of pre-molt carapace length (Table 2). The relationship between molting probability and pre-molt carapace length for European lobster was predicted from the relationship in Sørdalen *et al*. (2022) (reported in Supplemental Table 1). Therefore, for all growth calculations for European lobster, we combined the data on molt increment and molt frequency both in the protected area and the fished area to ensure our length-molt relationship estimates represented the widest range of size classes available.

### 2.3 Maximum Age

We assumed that individuals of both species do not live beyond 40 years of age. Historically, American lobsters have been observed to be as large as 217 mm carapace length or as much as 19 kg in weight, suggesting they can live for more than four decades (Lawton and Lavalli 1995). However, individuals of these larger sizes are extremely rare and difficult to capture (Cooper and Uzmann 1980; Sheehy *et al*. 1999). We used a maximum age of 40 years, given assumed rates of natural mortality and high fishing mortality in both American and Norwegian fisheries (ASMFC 2020; Fernández-Chacón *et al*. 2021).

### 2.4 Reproduction

Size-at-maturity and growth rate are dependent on sex in clawed lobsters. For both species, the cumulative probability a female was mature at a given carapace length was represented by logistic equations with parameters for size at 50% maturity (*L*_*mat*_) and shape parameters (*α* and *β* ). Values for each species are reported in Table 2. The functions used for this relationship, known as the maturation ogive for each species, were taken from the primary literature and differed slightly for each species. The equation for American lobster (ASMFC 2020) was:

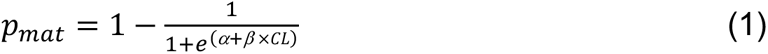

The other parameters are estimated from logistic regressions, which have been reported in slightly different functional forms. In American lobster, size at 50% maturity has shifted over time (Waller *et al*. 2019, 2021). Thus the most recent stock-wide estimates (and parameters) for Gulf of Maine female maturity were extracted from the 2020 American lobster stock assessment (see Appendix 1 in ASMFC 2020).

Since the size at 50% maturity (*L_mat_*) for the European lobster population in Southern Norway has not been determined, we relied on recent estimates of parameters for the size-specific probability of maturation obtained from Orkney in Scotland, which is similar in latitude to southern Norway (Coleman *et al*. 2023). The equation for European lobster (Walker 2005; Coleman *et al*. 2023) was:

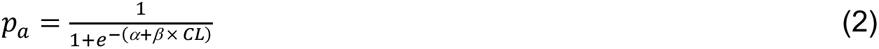

In American lobsters, males tend to reach physiological maturity (the ability to produce sperm and extrude spermatophores) at smaller sizes than females (Krouse 1973; Pugh *et al*. 2015). However, it is likely that males only participate in mating activities if they are at least the same size or larger than the mature females since females typically prefer larger males, and the male must be able to physically manipulate the female into mating position (Templeman 1933; Waddy and Aiken 1991; Sørdalen *et al*. 2018). Therefore, the functions used for female size-at-maturity were also used to approximate male functional maturity.

For both species, egg production in each year was calculated based on the proportion of mature females in the population in all age groups at a given time, their fecundity-at-size (estimated using functions in Table 2, with values reported in Supplemental Table 2), and their size-based spawning frequency. In year *t*, total egg production *E*_*t*_ is calculated as the summed egg production over all age classes *a*:

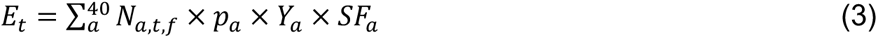

The total number of mature females in each age class in each year is represented by *N_a,t,f_*, where *f* signifies that we are only considering females. The probability a given female of age *a* is mature is *p*_*a*_ (given Eq. 1 and Eq. 2); this is multiplied by both the fecundity at size or age *Y* (here, given as a function of age, *Y_a_*) and the spawning frequency for a given age or size (*SF_a_*). This product is the number of eggs produced by a single female in each age class in a given year, which is then multiplied by the total number of females in that age class *N_a,t,f_*. The functions and parameters used to calculate *Y* and *SF* for each species are derived from empirical data and are given in Table 2. For American lobsters, we assumed female spawning frequency was every other year, as females alternate with molting years, until they are large enough to spawn twice per three years (Waddy and Aiken 1986; Waddy *et al*. 1995; ASMFC 2020). European lobsters tend to spawn every other year regardless of size, with some variability in the larger size classes (Sørdalen 2019).

The population in the model reaches a stable distribution of individuals given annual birth and death rates, sex, and average individual growth before fishing begins and the population reaches a new steady state. Recruitment was deterministic and was assumed to follow a density-dependent relationship between recruitment and spawning stock biomass. For both species, there is considerable uncertainty in stock recruitment relationships. for lobster (Sundelöf et al. 2015). Our approach to modeling stock dynamics is focused on the effects of fishing on adult age structure, not on recruitment overfishing, so we made the most conservative decision possible when selecting recruitment functions. For American lobster, a Ricker function is sometimes supported over a Beverton-Holt relationship (ASMFC 2020). However, preliminary analyses suggested our results were less sensitive to the choice of a Beverton-Holt recruitment function (Beverton and Holt 1957). The number of female recruits *R* in a year is given by

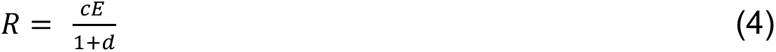

Total egg production *E* is calculated in Eq. 3. Parameter values (*c* and *d*) were arbitrary and are given in Table 2; we chose these parameters to have minimal influence on our results and conclusions after preliminary sensitivity analyses.

For European lobster, a Beverton-Holt stock-recruitment relationship has also been estimated (Bannister and Addison 1986). That relationship is given by

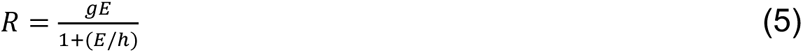

Again, specific parameter values (*g* and *h*) are given in Table 2. Individuals from both species are recruited to the model at 56 mm carapace length, which was assumed to correspond to roughly five years of age (ASMFC 2020), we only tracked age classes from five to 40, past which chances of survival were assumed to be negligible. Sex ratio at recruitment was assumed to be constant at 1:1.

### 2.5 Fishing Dynamics

As individuals recruit to the model, the simulated population will eventually reach an equilibrium size, depending on species- and sex-specific recruitment, growth, maturity, reproduction, and mortality rates (Table 2). The population dynamics of each age class over time are given by

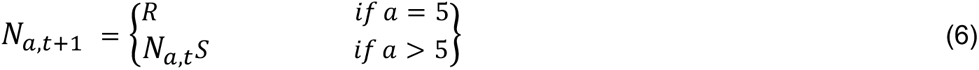

In Eq. 6, *N_a,t_* is the number of individuals of in each age class in year *t. S*urvival (*S*) to the next year is calculated according to the natural mortality (*M*) and, for age/size classes vulnerable to fishery, fishing mortality (*F*) (sex-specific values given in Table 2). Both mortality rates are instantaneous, such that *S* = *e^-(M+F)^*. Our simulated population grows from an arbitrary starting size at *t* = 0 until it reaches the unfished equilibrium, or steady state (Mangel 2006). After 100 years, we simulated size-selective fishing, with pressures ranging from lightly fished populations experiencing an annual harvest rate of approximately 25*t* (corresponding to an instantaneous fishing mortality rate (*F*) of 0.3) to intensely fished populations experiencing an annual harvest rate of approximately 60*t* (corresponding to an *F* of 0.9). We simulated the fished population for another 100 years, so that each simulation was for 200 years.

For each species, two types of ovigerous female protections were simulated to reflect actual conservation practices in the fisheries. A scenario without any female protections was also simulated for comparison. The first scenario of female protection was a 1-year protection for caught females, which represents a ban on harvest of egg-bearing females, consistent with Norwegian fishing practices. The second scenario of protection was a 4-year protection period representing the v-notch given to captured egg-bearing females in the Gulf of Maine. For both protection methods in the model, once caught and returned, females were removed from the fishable population for the relevant duration of time where they grew, reproduced, and survived until rejoining the fishable population. We assumed 100*t* fisher compliance with protection measures for egg-bearing females and size limits. For *H. gammarus*, sex-specific natural mortality was converted from the mean survival of lobsters in three protected areas in Norway for large and small individuals (Fernández-Chacón *et al*. 2021). Protected females and individuals outside of the species-specific slot limit therefore only experienced natural mortality and did not experience fishing mortality in our simulation.

Both American and European lobster fisheries have experienced high rates of harvest in the past and present; in the US lobster fishery, past stock assessments estimated *F* at greater than 1.0 (an annual harvest rate of 63*t*) in some years (ASMFC 2006, 2009; Kleiven *et al*. 2022). Simulating results for a wide range of fishing intensities provides insight on how exploitation interacts with different scenarios of female protection. For each species’ fishery, selectivity of the fishery is defined by a slot limit. Those individuals within the harvestable size range were subjected to fishing mortality in the model. Slot limits were different for each species and reflect current management practices (Table 1).

The model predicted the number of individuals of each sex at equilibrium, as well as their size-at-age with and without fishing. These values were used to quantify population size structure, size disparity between males and females, sex ratio, and egg production for each fishing intensity and management scenario. Size structure was calculated by summing the number of individuals by sex per size/age group for each scenario. Size disparity was calculated by subtracting the mean size of mature males from the mean size of mature females for each scenario. Sex ratio, or proportion female, was calculated by dividing the total mature female population by the total mature population of males and females. We also calculated annual egg production of all spawning females (year class strength) as the sum of all eggs that are produced by all spawning females in a given year under each fishing scenario, based on available fecundity-at-size estimates from the literature (Estrella and Cadrin 1995; Agnalt 2008).

## 3. RESULTS

### 3.1 Size structure

For both American and European lobsters, we used sex-specific growth functions to estimate size structure from age structure. In each species, males tended to grow faster than females (Figure 2). Although growth is continuous, for plotting purposes, we group individuals in 10 mm carapace length size bins. Males move through these bins more quickly than females of the same cohort (Figure 3). Differences between the two species, especially in the larger size classes, can be attributed to differences in the estimated rates of natural mortality and growth of each sex (Figure 3 A and C). To clearly see how fishing pressure and female protection measures affect population size structure, we compared two levels of fishing intensity representing moderate (*F* = 0.3) and intense (F = 0.9) fishing pressure (Figures 3-5). The predicted stable age distributions corresponding to all panels in Figures 3-5 can be seen in Supplemental Figures  1 and 2. Note the “intense” fishing scenario is still less than the level of *F* historically estimated for American lobsters (ASMFC 2006). The clearest effect of fishing is, as expected, reducing population abundance (numbers). Because our model includes strong assumptions about density-dependent recruitment (for which we have little empirical information) we are focused on relative trends and changes in demography, rather than predicting yield. The abundance of mature American lobster decreased by more than 60*t* with moderate fishing intensity (F = 0.3) in our model, depending on the female protection scenario. However, this level of fishing allowed for the persistence of both sexes into the size classes above the slot and even into the largest sizes of 156-165 mm carapace length (Figure 4). Under intense fishing (*F =* 0.9), the abundance of mature lobsters decreased by more than 88*t*, depending on the female protection scenario, and resulted in an extremely truncated size distribution for both sexes, with few adults persisting in the population above the slot limit. Management protecting females increased abundance for larger females under both levels of fishing, again more so in the 4-year protection scenario than the 1-year protection scenario (Figure 4).

**Figure 2.**
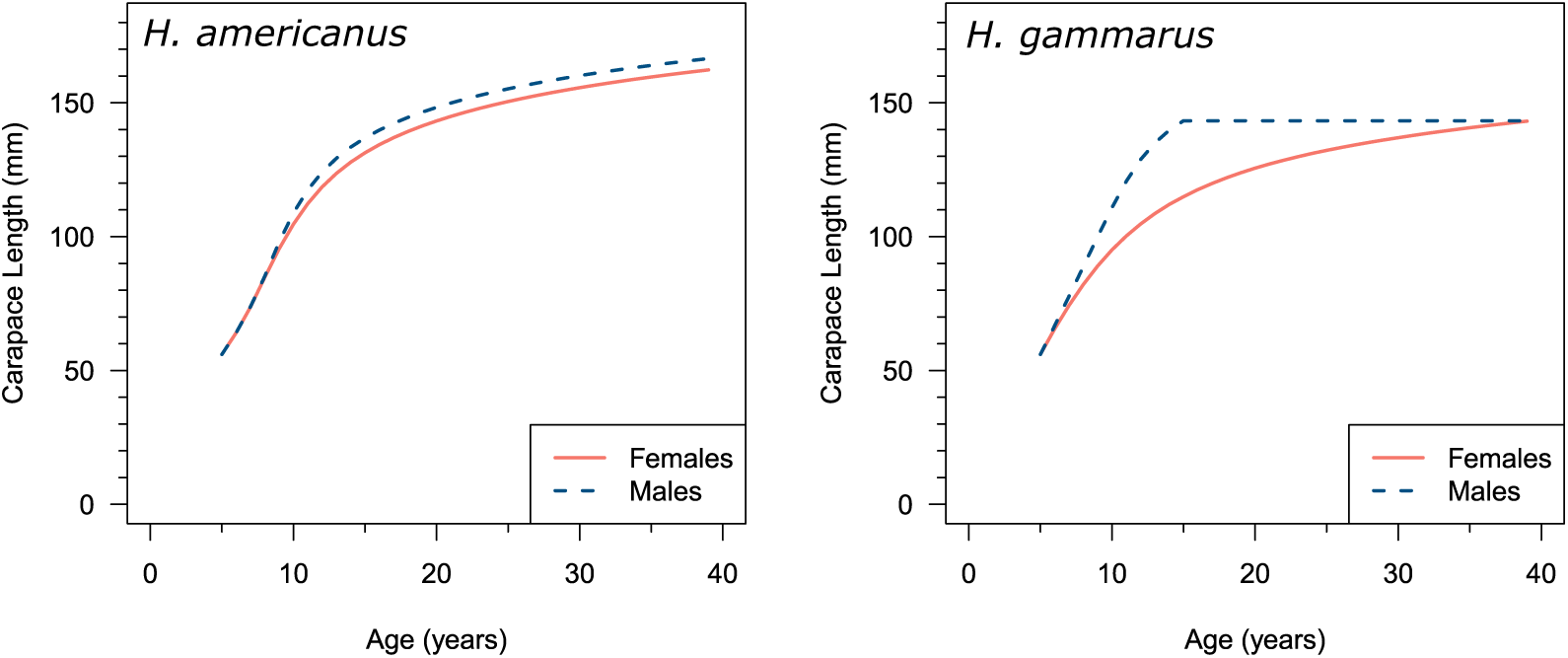
Predicted growth of American (*H. americanus*) and European (*H. gammarus*) lobsters over time. Age here is relative since these curves were generated using annual size-based probabilities of molting and molting increment for females and males of American (ASMFC 2009, 2020) and European (Sørdalen *et al*. 2022) lobsters.

**Figure 3.**
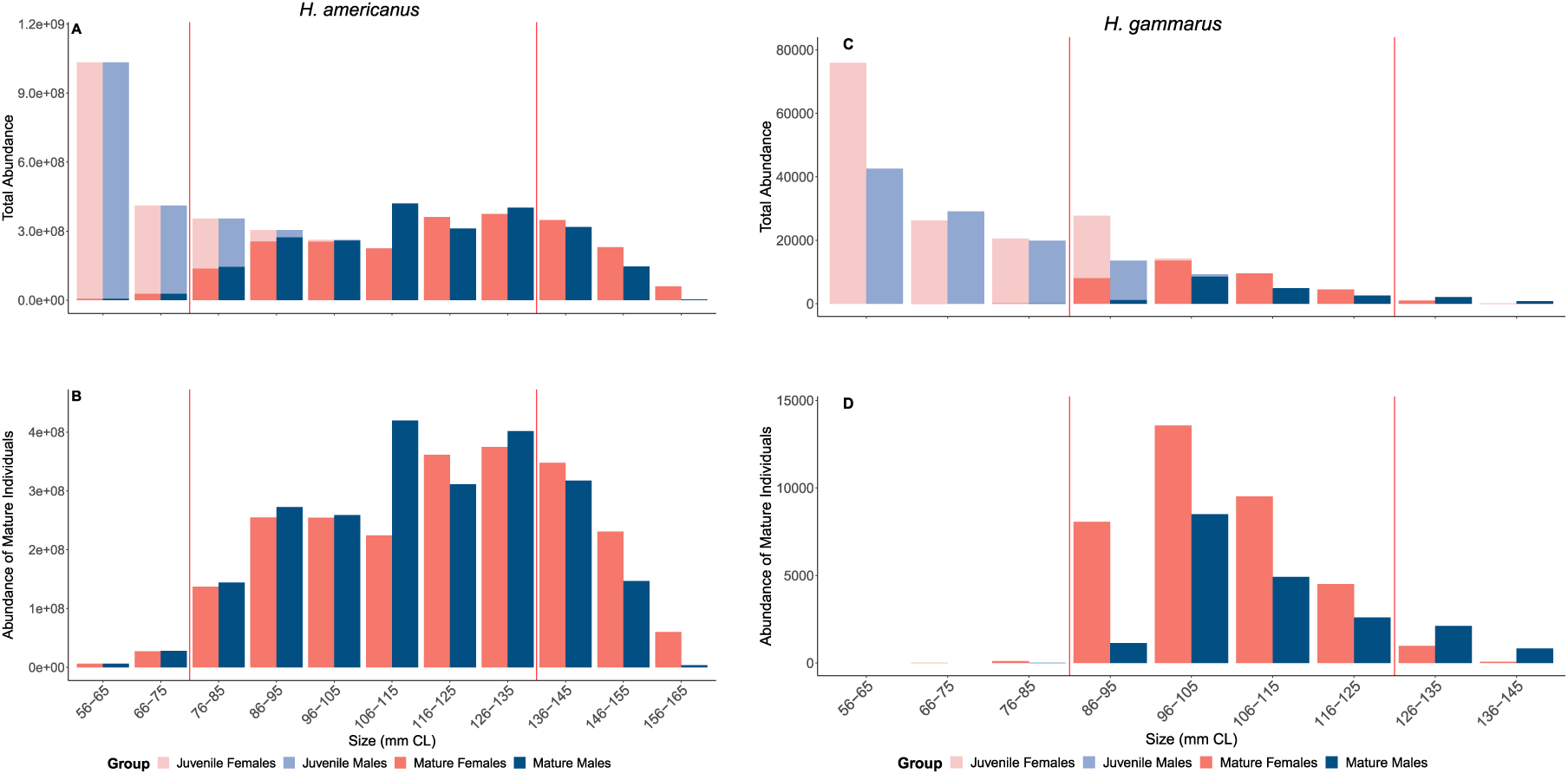
Size structure plots of the total population of American (*H. americanus*) in Panel A and European (*H. gammarus*) lobsters in Panel C over time without fishing. Size bins are in millimeters carapace length. Lighter versions of the colors for females and males indicate immature individuals, and darker colors are mature. Panels B and D show only the mature population for American and European lobsters, respectively. The size bins in between the vertical red lines contain the harvestable size limits for reference. The American lobster slot limit is 83-127 mm carapace length. The slot limits for European lobster are 86.9-114.76 mm (females) and 88.6-117.09 mm (males) carapace length.

**Figure 4.**
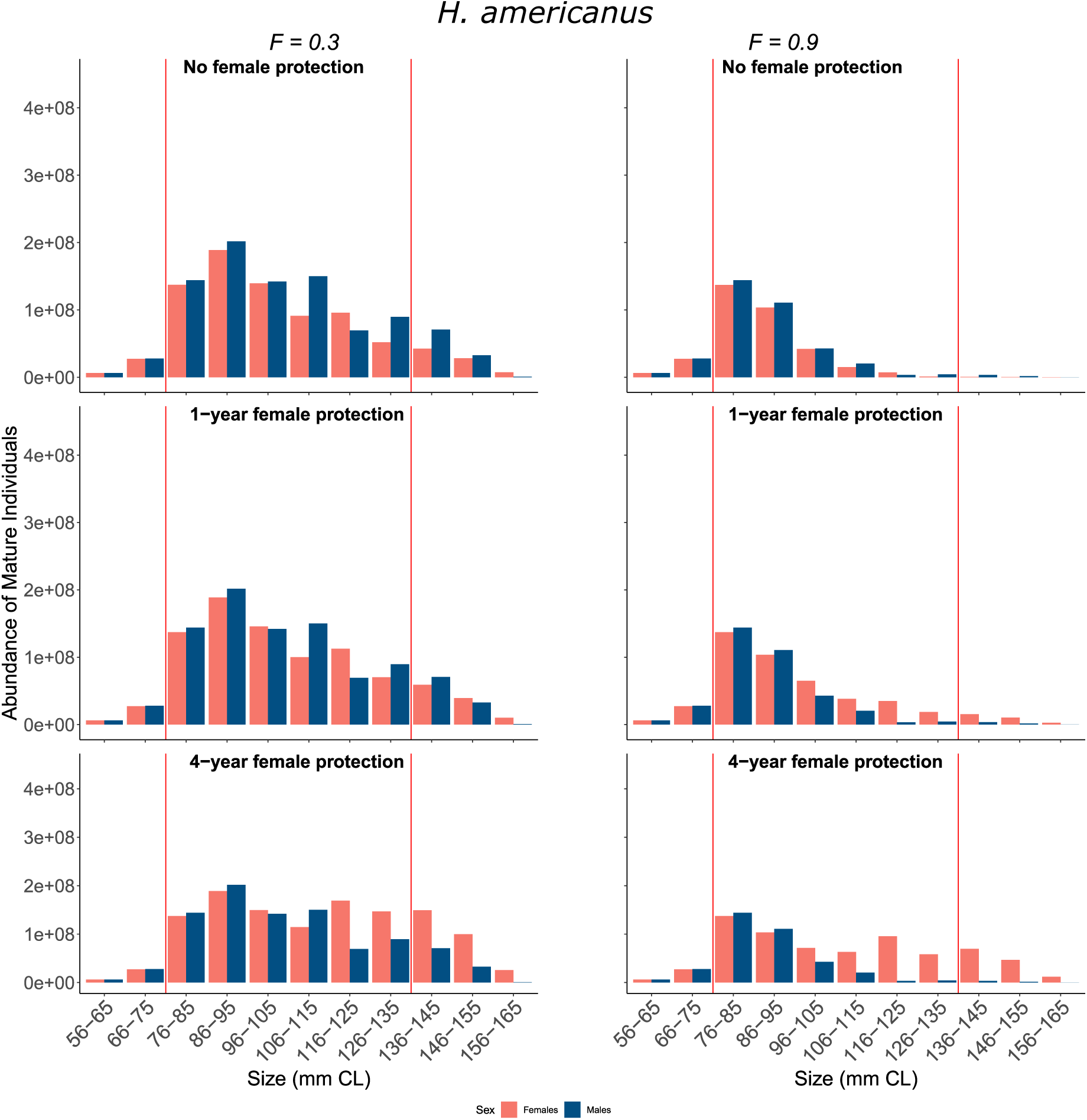
Size structures of American lobsters with and without female protections during fishing compared to the unfished size structure. Size bins are in millimeters carapace length. Note that these are for mature individuals only at *F* = 0.3 and *F* = 0.9. The first row shows size structures after fishing without any female protections. The second row shows size structures after fishing with 1-year protections for ovigerous females. The third row shows after fishing with 4-year protections for ovigerous females. The size bins in between the vertical red lines contain the harvestable size limits of 83-127 mm carapace length for American lobster.

**Figure 5.**
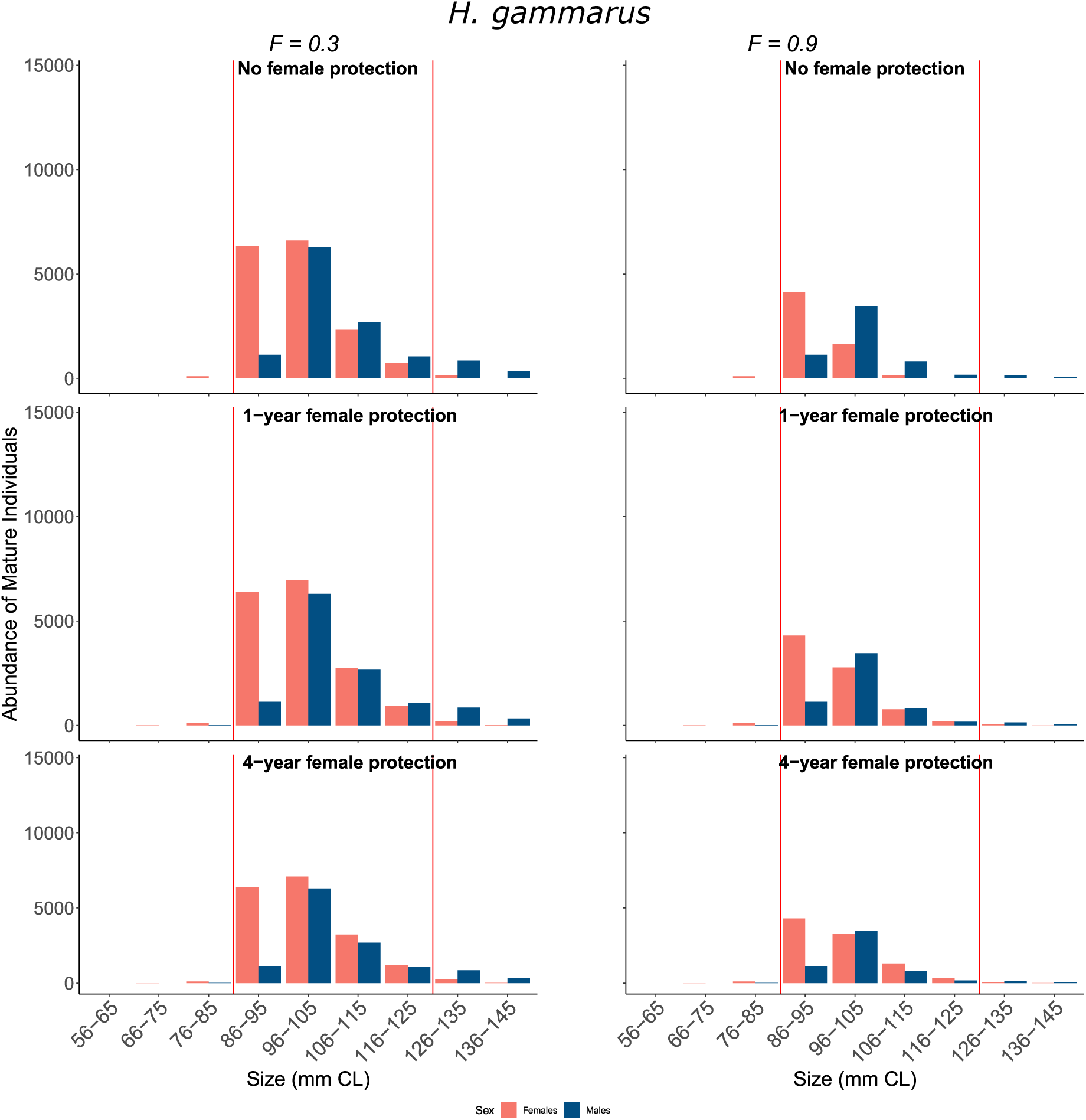
Size structures of European lobsters with and without female protections during fishing compared to the unfished size structure. Size bins are in millimeters carapace length. Note that these are for mature individuals only at F = 0.3 and F = 0.9. The first row shows size structures after fishing without any female protections. The second row shows size structures after fishing with 1-year protections for ovigerous females. The third row shows after fishing with 4-year protections for ovigerous females. The slot limits for European lobster are 86.9-114.76 mm (female) and 88.6-117.09 mm (males) carapace length.

Trends from our model for European lobster showed that moderate fishing pressure (*F* = 0.3) reduced abundance for all size classes above the minimum legal size, but relative declines in abundance were less dramatic than for American lobster, ranging from 36-39*t* depending on the scenario of female protection. Moderate fishing allowed for adults of both sexes to survive long enough to outgrow the slot and reach the protected larger size classes (Figure 5). In model scenarios with female protections, large females above the slot limit were slightly more abundant (discussed below). Under intense fishing pressure (*F* = 0.9), numbers of mature lobsters decreased by 60-63*t* across all female protection scenarios. The size structure was more truncated, especially for males, in the largest harvestable size classes, with very few individuals of either sex surviving to grow larger than the slot. Even with the 4-year protection, in our model of European lobster, there were very few females in the size bin just above the slot limit compared to the unfished population (Figure 5).

### 3.2 Size Disparity Between Sexes

The size distribution of populations of American lobsters responded differently than European lobsters to the same scenarios of female protections and range of fishing mortality, due in part to inherent differences in the sizes of mature individuals evident in the unfished scenario (Figure 3). This resulted in species differences in the size disparity between sexes (Figure 6). Without fishing, American lobster mature males were on average 4 mm carapace length (CL), or about 3*t*, larger than mature females (Figure 6). Without specific measures protecting females during fishing, males remained larger than females, by an average of 3 to 7 mm, across fishing levels. With one year of protection, females were up to 5 mm (CL) smaller than males until a fishing level of 0.6, where males and females were on average the same size. Above *F* = 0.6, the mean difference increased so that females were up to 5 mm (CL) larger than males at *F* = 0.9. Management measures protecting females for four years led to the most extreme size disparity, where females were on average 2.5 mm CL larger than males at *F* = 0.3, but as much as 8 to 16 mm larger than males at *F* ≥ 0.5. In other words, at *F* ≥ 0.5 the average male was 7 to 14*t* smaller than the average female. This predicted pattern is consistent with the size disparity observed in the Georges Bank sub-stock of American lobster (ASMFC 2020; T. Pugh, *personal observation*).

**Figure 6.**
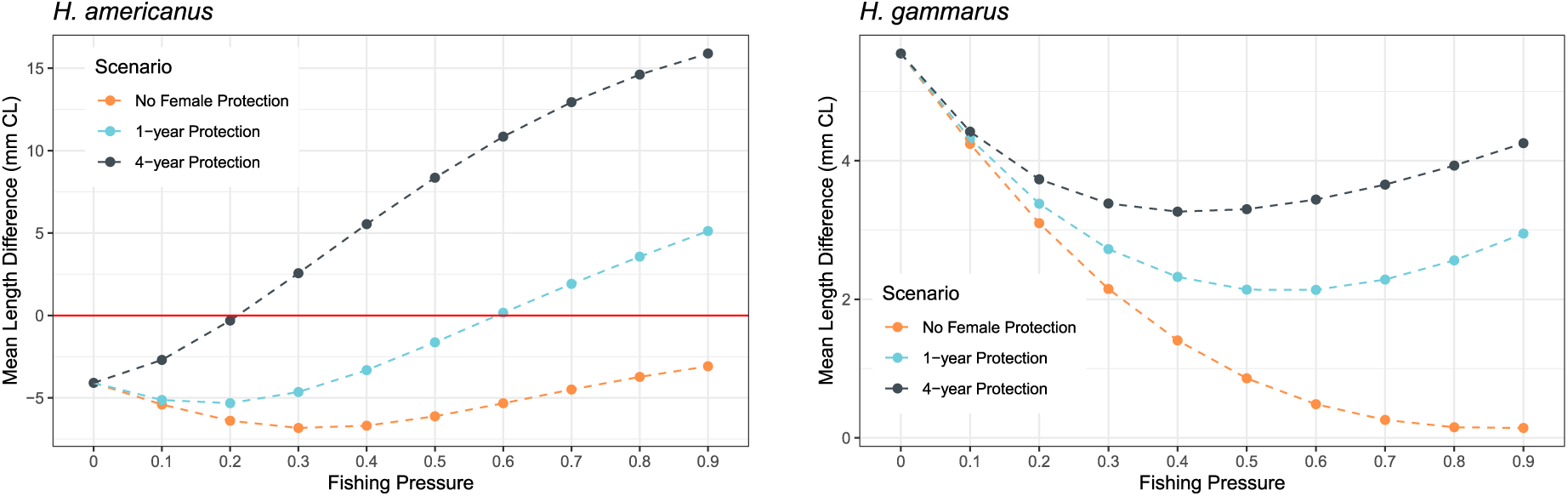
Mean size disparity by sex calculated by subtracting mean mature male carapace length from mean mature female carapace length for American (*H. americanus*) and European (*H. gammarus*) lobsters. The red horizontal lines indicate the threshold (zero) below which males are larger than females on average, or above which females are on average larger than males.

In all scenarios, the predicted average size of European lobster females was larger than males, because the low survival of males led to few in the largest size classes, even in the unfished population. Therefore, without fishing, the size difference between European lobster males and females was predicted to be opposite to American lobster, with mature females on average 5.5 mm (CL) or 4*t* larger than males (Figure 6). This size disparity in favor of females decreased slightly with low to moderate fishing pressure (*F* = 0.1 – 0.4) in all female protection scenarios. With *F* > 0.4, females remained larger than males, particularly so in the 4-year protection scenario. Only in the scenario without female protection measures at very high fishing pressure (*F* > 0.7), did females and males show very little size disparity (less than 2*t* difference in CL). The greatest disparity occurred in the 4-year protection scenario with intense fishing pressure (*F* > 0.8) where females were up to 4*t* larger than males (4.5 mm in CL).

### 3.3 Sex Ratio

For American lobster, sex ratios were fairly even (near 1:1) without female protections, but became more biased toward females as protection measures increased (from 1 to 4 years) and as fishing intensity increased (Figure 7). In the 4-year protection scenario, the bias of the sex ratio toward females reached 65*t* female at very high fishing pressure (*F* = 0.9). For European lobster, as with the pattern of size disparity, the unfished population was biased toward females. This female bias in sex ratio persisted across levels of fishing in every female protection scenario and fishing pressure, with very little difference arising from female protection scenarios. The model predicted female European lobsters consistently comprised between 56-70*t* of the mature population, such that 70*t* of the mature population was female without fishing, which is consistent with patterns observed in unfished areas (Fernández-Chacón et al. 2021). At low fishing pressure (*F* = 0.1 – 0.3) the proportion of females in the population for both the 1-year and 4-year protection scenarios remained very similar to the scenario without female protections. As fishing intensity increased, the proportion female in the 1-year protection scenario remained within 0.03*t* of those in the 4-year protection scenario. At *F* ≥ 0.5, females without any protections decreased to between 56 – 59*t* of the total population.

**Figure 7.**
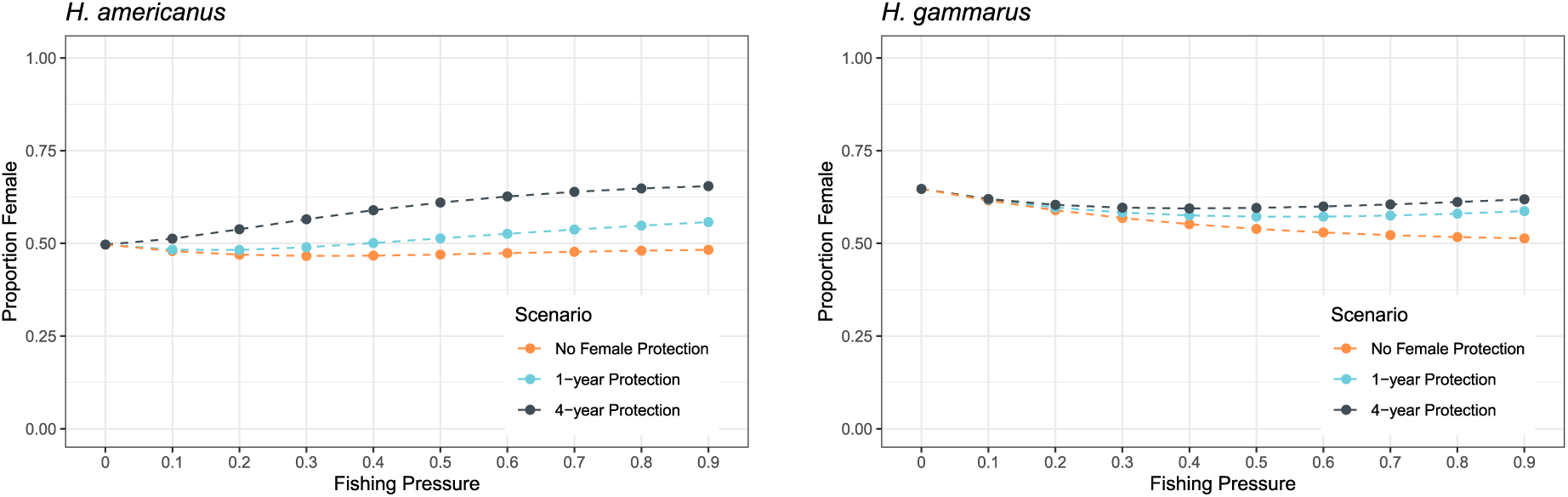
Proportion female for American (*H. americanus*) and European (*H. gammarus*) lobsters over in three alternative female protection scenarios and varied levels of fishing pressure.

### 3.4 Egg Production

As egg production is a function of the number of spawning females in the population, even the lowest fishing pressure had a negative effect on total egg production for both species (Figure 8). Across all scenarios for both species, increasing fishing intensity reduced egg production compared to unfished production levels, with the largest declines occurring as *F* increased from 0 to around 0.4. Of the two female protection scenarios, the 4-year protection resulted in the least decline in egg production, especially at the highest levels of fishing pressure. The effectiveness of the two measures of female protection in increasing egg production were most evident for moderate to intense levels of fishing, particularly for American lobsters.

**Figure 8.**
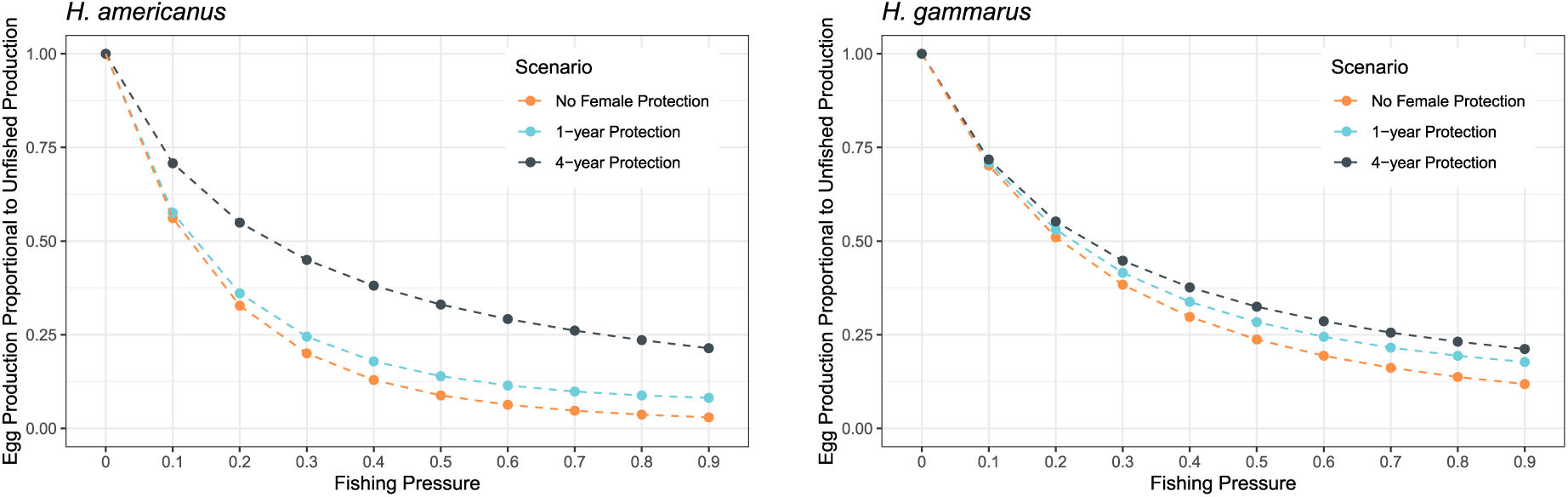
Egg production during fishing scaled to the unfished level of egg production for American (*H. americanus*) and European (*H. gammarus*) lobsters in three alternative female protection scenarios and varied levels of fishing pressure.

Overall, the egg production in American lobster was more sensitive to changes in fishing pressure compared to the European lobster, meaning that the *relative* impact of fishing on egg production was greater in American lobster. While American lobster population numbers were overall much higher than the European lobster, egg production declined to a larger extent with fishing in American lobster (Figure 8). With increasing levels of fishing pressure, egg production declined from 71*t* to 4*t* of unfished levels for American lobster. When fishing pressure exceeded *F* = 0.6 without any female protections, egg production was extremely low. Although the 1-year female protection improved egg production slightly, the 4-year protection consistently led to up to 13*t* higher egg production at all levels of fishing pressure.

In European lobster, the sensitivity of egg production to changes in fishing pressure was slightly lower compared to American lobster. Across all management scenarios, egg production with increasing fishing pressure declined from 70*t* to 12*t* of unfished production levels. At low fishing pressure, both forms of female protections had minimal impact on egg production, but their efficacy in improving annual egg production became more apparent as fishing pressure approached moderate levels (Figure 8), with protection of females for four years being more effective than one year of protection.

## 4. DISCUSSION

The aim of this study was to assess the effects of different management regulations, particularly two types of protection applied to reproductive females, on population size structure, sex ratio, and egg production in response to varying levels of fishing intensity for both American and European lobster fisheries. As expected, fishing led to large declines in population size and commensurate decreases in egg production, which occurred even with slot limits in place under intense fishing. Implementing one year of protection for egg-bearing females allowed some females to reach larger sizes, resulting in improved egg production in both species, particularly at higher fishing pressure, consistent with other research (Sundelöf *et al*. 2015). The 4-year protection scenario further enhanced egg production, most notably for American lobsters. However, particularly for American lobsters, female protections also led to skewed sex ratios and increased size differences between females and males, which could potentially have unintended consequences for population productivity if the disparity between males and females affects fertilization success. Differences in the size distributions of American and European lobster under different fishing scenarios in Figures 4 and 5 are due to the high rates of both growth and mortality of males estimated from experimentally protected and fished populations of European lobster (Sørdalen *et al*. 2022). These differences in life history traits led to less impactful effects of female protection measures, especially 4-year protection, on our modeled populations of European lobster.

### Effects of fishing on productivity

The nuanced implications of fishing mortality on population size are often overlooked in discussions surrounding resource management focused on yields. We show that realistically intense levels of fishing pressure led to a substantial reduction in abundance, and that even low levels of fishing had serious effects on annual egg production (spawning potential) in both species. Our results suggest that management will have to drastically reduce *F* in order to see signs of recovery. Low population biomass and spawning potential can profoundly affect a population’s ability to recover from overfishing or to maintain resilience to natural or anthropogenic environmental variation. Depleted populations may reach a critical threshold of density, where decreased fitness of the remaining individuals may negatively affect population growth rate, a phenomenon known as depensation (Allee 1938; Hutchings 2014). One mechanism contributing to depensation is when individuals are unable to reproduce successfully because they are unable to locate a viable mate (Errington 1940; Gascoigne et al. 2009). Variable reproductive success in the Caribbean spotted spiny lobster (*P. guttatus*) in isolated patch reefs has been attributed to lack of mating opportunities and subsequently population-level depensation (Robertson and Butler 2009). Depleted populations also yield less over longer timescales, resulting in a lose-lose situation both from a perspective of ecosystem function and from a sustainable management view. An additional factor undermining population productivity comes from the fact that fisheries consistently remove the largest individuals, which can inadvertently lead to selection for slower growing individuals, smaller body size, and smaller size at maturity (Hutchings and Rowe 2008). In other words, lower population sizes not only diminish yield to the fisheries at present but can also further diminish returns by eroding genetic variation in growth that may be important for long-term sustainable yield (Conover and Munch 2002). Both clawed lobster species studied here show both temporal and spatial variation in the size at maturation, which could reflect the evolutionary effects of harvest, in addition to recent increases in temperature, which is known to lead to earlier maturation (Le Bris et al. 2017; Waller et al. 2021).

### Effects of fishing on population size structure

We were able to link population age structure to population size structure by calculating the expected growth trajectory of the average individual in a deterministic environment for each species. We incorporated discontinuous growth by calculating mean size at age given annual growth probabilities, which opens new possibilities for crustacean modelling. For simplicity, we did not incorporate environmental variability or density effects on growth, instead relying on parameters and equations that have been estimated previously through field and lab studies. Our results illustrate how fishing reduces population abundance, erodes size structure and spawning potential, and may lead to the evolution of faster growth and smaller sizes at maturity. These effects could explain the worrisome status of the European lobster populations in Norway, which have declined by 92*t* over the past 90 years (Kleiven *et al*. 2022). While American lobster in the Gulf of Maine is still at very high abundance, the Southern New England US lobster stock is significantly depleted and may be at or near critical thresholds (ASMFC 2020; Pugh *et al*. 2023).

Removal of large females from the population can negatively affect overall productivity and population resilience as large females in aquatic environments are not only more fecund, but can exhibit different spawning behavior to maximize reproductive success (a bet-hedging strategy) (Hixon *et al*. 2014). Age-truncated populations have also been found to track environmental fluctuations more closely, affecting the resilience of the populations, and consequently destabilizing the fisheries (Wright and Trippel 2009; Rouyer *et al*. 2011). This effect of fishing on population demography is exaggerated at higher intensities of fishing, which was clearly demonstrated in our results for both species (Figures 4 and 5), even though both simulated fisheries have a slot limit (minimum and maximum harvest size). Management measures taken to protect reproductive females are designed to preserve size-based fecundity, thereby promoting the survival of larger, more fecund females to support overall population growth. However, our model suggested protecting only females within the harvestable slot resulted in females surviving longer and thus outgrowing their male counterparts.

The species trends in size-disparity across fishing levels arise (Figure 6) from these differences in estimates of growth and natural mortality. The disparity in average size of American lobsters was biased toward larger males, but this relationship reversed with fishing, especially in scenarios with protection of mature females (Figure 6). For European lobster, in all model scenarios, on average, females were larger than males (even though males grow much faster; Figure 2) because of the low survival of males. This disparity initially decreases as fishing pressure increases, as both large females and males are removed from the population.

However, the female-biased disparity between the sexes increased slightly at higher fishing in scenarios with female protections (Figure 6). This is somewhat in line with observations from southern Norway where females on average were larger than males in no-take areas, (where fishing mortality is close to zero), a size disparity that was slightly amplified in fished areas (Sørdalen *et al*. 2020, Table 1). However, these trap-based studies have also been found to under-sample large males, potentially biasing the growth and survival parameters used here (Helms 2023).

### Effects of fishing on the size disparity between sexes

The truncation of the adult size distribution and changes to sex ratios in response to fishing and female protections (Figure 7) can be expected to influence the mating system of both American and European lobsters, considering their similar mating behavior. Both depletion of large individuals and skewed sex ratios in larger size classes can limit the opportunity to find mates and affect the opportunity for and the strength of sexual selection (Kokko *et al*. 2012). Female lobsters typically select a male with large body size and large crusher claw, which indicate dominance status (Talbot and Helluy 1995; Atema and Steinbach 2007; Sørdalen *et al*. 2018). In European lobster, size-selective fishing appears to disrupt these sexually selected traits, suggesting that fishing can alter mating behavior and weaken sexual selection on male traits (Sørdalen *et al*. 2018, 2020). Although females may mate with smaller males when faced with scarcity of choice, experimental studies suggest that smaller males may not provide sufficient sperm to fertilize the clutch of larger females, potentially leading to sperm limitation (Gosselin *et al*. 2003; Pugh 2014; Pugh *et al*. 2015; Tang *et al*. 2019). While some evidence from other exploited crustacean populations suggests that changes in population sex ratio and size structure may result in sperm limitation (Pardo *et al*. 2015; Baker *et al*. 2022), this relationship still needs further investigation for clawed lobsters. The results of our model are sensitive to sex- and size-based mortality, and do not address potential variation in mating success or fertilization rates resulting from predicted size-disparities and sex-ratios. Further understanding of these relationships in clawed lobsters is needed to fully understand the potential for sperm limitation to limit their population productivity beyond direct effects of reduced egg protection.

### Effects of fishing on spawning potential

The spawning potential (egg production) of each species declined dramatically with fishing intensity, but was affected differently by each measure of female protection (Figure 8). In this study, European lobsters did not experience the same degree of improvement in egg production or size disparity as American lobsters with female protections. The model outputs show that v-notching as a form of 4-year protection is an effective means of boosting egg production if the female can survive long enough to benefit from this. Because of the assumed low natural mortality of American lobster, females were able to survive through the 4 years of protection to continue growing and reproducing. In species with higher estimates of natural mortality rates, including the European lobster, v-notching (represented here as 4-year protection) is predicted to only marginally enhance egg producing potential compared to a ban on harvest of egg-bearing females alone (Sundelöf et al. 2015). Future studies could test the hypothesis that European lobster would benefit from a narrower slot limit (a lower maximum legal size) to improve the chances both sexes outgrow the size classes vulnerable to harvest. However, even with female protections, our results show that fishing at high intensities may lower spawning potential enough to reduce the overall population’s resilience to fishing or poor environmental conditions. Intensively fishing a population to a spawning potential ratio of less than 0.4 has been suggested to be a critical threshold past which populations do not have a sufficient spawning potential to buffer natural variability in recruitment (Wiedenmann et al. 2006; Zhou et al. 2020; Conrad and Kindsvater 2024). In Figure 8, our model predicts this threshold occurs at low to moderate levels of fishing, depending on the species and female protection scenario.

### Survival and growth rates

The literature-based estimates of survival and growth rates that we used contribute to our results, including the effects of slot limits and protections such as v-notching on size structure and egg production, because these rates determine the time an individual spends in the size classes vulnerable to fishing and the probability of surviving (Chong-Montenegro and Kindsvater 2022). We based survival on mortality estimates for European lobster calculated from a trap-based mark-recapture study (Fernández-Chacón *et al*. 2021), and like any study conducted in this manner, estimates are dependent on season, sex and size-dependent catchability, trap design (Miller 1990), and assumptions around immigration/emigration (Skerritt *et al*. 2023). By contrast, the natural mortality parameter used for American lobster was the value used by recent stock assessments for Gulf of Maine lobsters; it does not differentiate by sex and is not empirically derived (ASMFC 2020). If this parameter is an underestimate of the natural mortality of American lobster, then we may be overestimating the effectiveness of v-notching and results may be more similar to the modeled European lobster results currently shown (and vice versa). Future studies to simulate both possibilities would clarify this discrepancy, and improved empirical data on natural mortality is clearly desirable.

The growth parameters used in this study for European lobster are based on recent tag-recapture data that estimated molting probability and molt increment (Sørdalen *et al*. 2022). We combined data from protected and fished areas to maximize the size range used in the parameter estimates. This choice may have impacted some of the modeled differences between male and female growth since lobsters from fished areas do not grow as fast as those subjected to protection, an effect particularly evident in females (Sørdalen et al. 2022). The plateauing of male growth (see Figure 2) is likely due to the combined effect of insufficient empirical data to estimate molt parameters at the larger sizes and the relatively high natural mortality estimate used for males. The growth parameters for American lobster are mainly from older data that have been recycled over a number of stock assessments (ASMFC 2020). This could be an indication that updated estimates of natural mortality, growth, and fecundity are necessary to accurately represent American lobster population dynamics into the future, especially since it has already been shown that their size-at-maturity has been changing over time (Waller *et al*. 2019).

The comparative nature of our work demonstrates the importance of the estimated natural mortality and other life history parameters to modeled results, and shows that accurate knowledge of species life histories is needed for improved modelling and for truly understanding the effect of fishing on population demography and dynamics. Yet intense fishing activities in the past decades have left few areas unaffected by fishing, thereby limiting the availability of reference areas for collecting data on life history parameters and natural behavior. No-take areas have been adopted as management tool in recent years, with 60 such areas already implemented across southern Norway, aimed at aiding the recovery of local populations of European lobster (Moland *et al*. 2021; Knutsen *et al*. 2022). Remarkably, these Norwegian protected areas have demonstrated that they need not be large to elicit population-level responses, likely in part due to the coastal fjord landscape and the high site fidelity of individual European lobsters (Moland et al. 2021). The observed changes in demographic structures, phenotypic composition, mating behavior, and reproductive output within reserves in Norway and other locations have provided valuable insights into the effects of fishing and the impact of release from fishing pressure (Bertelsen and Matthews 2001; Goni *et al*. 2003; Sørdalen *et al*. 2018, 2020, 2022; Fernández-Chacón *et al*. 2021). Our simulation results clearly showed differences in the demographics between the unfished population (comparable to a reserve), very low levels of *F*, and higher levels of fishing intensity. Whether no-take areas as implemented are supporting overall population growth and the fishery through spillover is yet to be determined.

## 5. CONCLUSION

The sensitivity of overall population size and size structure to fishing pressure as shown in our results for both species emphasizes the need for implementing targeted measures of controlling fishing mortality, to prevent further depletion and facilitate recovery of lobster populations. Such measures could range from narrowing the harvest window of minimum and maximum size limits, or more directly controlling *F* by implementing catch quotas. Understanding how management decisions influence population structure is crucial, particularly in light of the predicted decline in American lobster recruitment and that there are still no signs of recovery for European lobster (Sundelöf et al. 2015; Le Bris *et al*. 2018; Oppenheim *et al*. 2019; Kleiven *et al*. 2022). As managers explore options for addressing the challenges faced by these two species, the development and refinement of models like the one we have presented here can serve as critical tools for evaluating management strategies and forecasting possible outcomes.

## Author Contributions

Conceptualization: K.T., T.K.S., T.L.P, H.K.K., Methodology and Software: K.T., H.K.K., Writing original draft: K.T., Reviewing and Editing: K.T., T.K.S., T.L.P., H.K.K.

## Acknowledgements

K.T. and H.K.K. were supported by the Department of Fish and Wildlife Conservation at Virginia Tech. This study was completed as part of a K.T.’s Master of Science thesis, and she would like to thank her committee, particularly Leandro Castello. We would also like to thank Kim Halvorsen for assistance with growth data and for insightful conversations about *H. gammarus*. K.T. also thanks Hailey Conrad for assistance troubleshooting R code.

---

**Supplemental Table 1.** Probabilities of molting given pre-molt carapace length (CL) for females and males of European lobster (*H. gammarus*). Data from Sørdalen *et al*. (2022) on lobsters in fished and protected areas were combined, then converted from total length to carapace length for this study.

| Pre-molt CL | Probability Molting |  |  |  |  |  |  |  |
| --- | --- | --- | --- | --- | --- | --- | --- | --- |
|  | Female | Male |  |  |  |  |  |  |
| 56 | 0.954438 | 0.998032 | 93 | 0.70734 | 0.955662 | 133 | 0.190929 | 0.417955 |
| 57 | 0.951875 | 0.997854 | 94 | 0.693841 | 0.951792 | 134 | 0.181417 | 0.395729 |
| 58 | 0.949056 | 0.997675 | 95 | 0.6823 | 0.947606 | 135 | 0.172254 | 0.378258 |
| 59 | 0.945928 | 0.997459 | 96 | 0.669446 | 0.943471 | 136 | 0.165199 | 0.356878 |
| 60 | 0.942785 | 0.99723 | 97 | 0.656279 | 0.939029 | 137 | 0.156721 | 0.336017 |
| 61 | 0.93953 | 0.996999 | 98 | 0.644049 | 0.933856 | 138 | 0.148578 | 0.319787 |
| 62 | 0.9361 | 0.996749 | 99 | 0.630305 | 0.928179 | 139 | 0.142326 | 0.300124 |
| 63 | 0.932807 | 0.99645 | 100 | 0.616654 | 0.922635 | 140 | 0.134833 | 0.281124 |
| 64 | 0.929357 | 0.996128 | 101 | 0.602488 | 0.91617 | 141 | 0.127654 | 0.266481 |
| 65 | 0.925385 | 0.995805 | 102 | 0.587108 | 0.90915 | 142 | 0.122157 | 0.255807 |
| 66 | 0.921234 | 0.995418 | 103 | 0.572675 | 0.902291 |  |  |  |
| 67 | 0.916776 | 0.995005 | 104 | 0.559467 | 0.894289 |  |  |  |
| 68 | 0.911843 | 0.994589 | 105 | 0.546175 | 0.885699 |  |  |  |
| 69 | 0.906924 | 0.994089 | 106 | 0.531627 | 0.87731 |  |  |  |
| 70 | 0.902283 | 0.993558 | 107 | 0.516899 | 0.867523 |  |  |  |
| 71 | 0.896988 | 0.993023 | 108 | 0.503462 | 0.857146 |  |  |  |
| 72 | 0.891413 | 0.992377 | 109 | 0.488632 | 0.847024 |  |  |  |
| 73 | 0.886097 | 0.99163 | 110 | 0.47273 | 0.835228 |  |  |  |
| 74 | 0.879958 | 0.990856 | 111 | 0.457912 | 0.822887 |  |  |  |
| 75 | 0.873675 | 0.990035 | 112 | 0.443497 | 0.810875 |  |  |  |
| 76 | 0.866951 | 0.989122 | 113 | 0.428896 | 0.796905 |  |  |  |
| 77 | 0.859425 | 0.988222 | 114 | 0.416458 | 0.782497 |  |  |  |
| 78 | 0.852112 | 0.987164 | 115 | 0.401268 | 0.768517 |  |  |  |
| 79 | 0.845206 | 0.985996 | 116 | 0.386254 | 0.753918 |  |  |  |
| 80 | 0.837312 | 0.984842 | 117 | 0.374399 | 0.737259 |  |  |  |
| 81 | 0.829186 | 0.983594 | 118 | 0.359801 | 0.71995 |  |  |  |
| 82 | 0.821436 | 0.982141 | 119 | 0.34544 | 0.703471 |  |  |  |
| 83 | 0.812478 | 0.98051 | 120 | 0.334156 | 0.684753 |  |  |  |
| 84 | 0.802719 | 0.978914 | 121 | 0.320325 | 0.665627 |  |  |  |
| 85 | 0.793065 | 0.977048 | 122 | 0.306783 | 0.64964 |  |  |  |
| 86 | 0.784099 | 0.974972 | 123 | 0.296191 | 0.626156 |  |  |  |
| 87 | 0.774081 | 0.972935 | 124 | 0.283265 | 0.604382 |  |  |  |
| 88 | 0.763528 | 0.970546 | 125 | 0.270663 | 0.586642 |  |  |  |
| 89 | 0.753681 | 0.967912 | 126 | 0.260849 | 0.564143 |  |  |  |
| 90 | 0.742565 | 0.965321 | 127 | 0.251267 | 0.541395 |  |  |  |
| 91 | 0.731203 | 0.962273 | 128 | 0.239636 | 0.52306 |  |  |  |
| 92 | 0.720504 | 0.958945 | 129 | 0.228355 | 0.500058 |  |  |  |
|  |  |  | 130 | 0.219613 | 0.477057 |  |  |  |
|  |  |  | 131 | 0.20904 | 0.458721 |  |  |  |
|  |  |  | 132 | 0.198821 | 0.440496 |  |  |  |

**Supplemental Table 2.** Fecundity-at-size estimates for American lobster (*H. americanus*; Estrella and Cadrin 1995) and European lobster (*H. gammarus*; Agnalt 2008). Functions and parameters are in Table 2.

| *H. americanus* |  | *H. gammarus* |  |
| --- | --- | --- | --- |
| Carapace Length (mm) | Average fecundity | Carapace Length (mm) | Average fecundity |
| 73.72 | 295 | 74.30 | 1781 |
| 84.84 | 8117 | 82.13 | 5445 |
| 95.44 | 12374 | 89.03 | 8679 |
| 104.66 | 17213 | 95.04 | 11493 |
| 112.32 | 22166 | 100.23 | 13920 |
| 118.54 | 26891 | 104.71 | 16018 |
| 123.61 | 31241 | 108.63 | 17853 |
| 127.80 | 35198 | 112.02 | 19443 |
| 131.32 | 38796 | 114.92 | 20798 |
| 134.34 | 42083 | 117.56 | 22033 |
| 136.97 | 45103 | 119.87 | 23115 |
| 139.28 | 47895 | 121.97 | 24099 |
| 141.35 | 50491 | 123.86 | 24985 |
| 143.22 | 52917 | 125.58 | 25788 |
| 144.91 | 55196 | 127.13 | 26515 |
| 146.47 | 57346 | 128.54 | 27172 |
| 147.90 | 59381 | 129.86 | 27790 |
| 149.23 | 61314 | 131.11 | 28377 |
| 150.47 | 63156 | 132.23 | 28901 |
| 151.63 | 64916 | 133.29 | 29398 |
| 152.72 | 66601 | 134.29 | 29865 |
| 153.75 | 68218 | 135.23 | 30304 |
| 154.72 | 69773 | 136.12 | 30721 |
| 155.64 | 71272 | 136.95 | 31113 |
| 156.51 | 72717 | 137.78 | 31500 |
| 157.35 | 74115 | 138.56 | 31865 |
| 158.15 | 75467 | 139.30 | 32211 |
| 158.91 | 76777 | 139.99 | 32536 |
| 159.64 | 78049 | 140.68 | 32858 |
| 160.34 | 79284 | 141.33 | 33161 |
| 161.01 | 80484 | 141.94 | 33449 |
| 161.66 | 81653 | 142.55 | 33735 |
| 162.29 | 82791 |  |  |

**Supplemental Figure 1.**
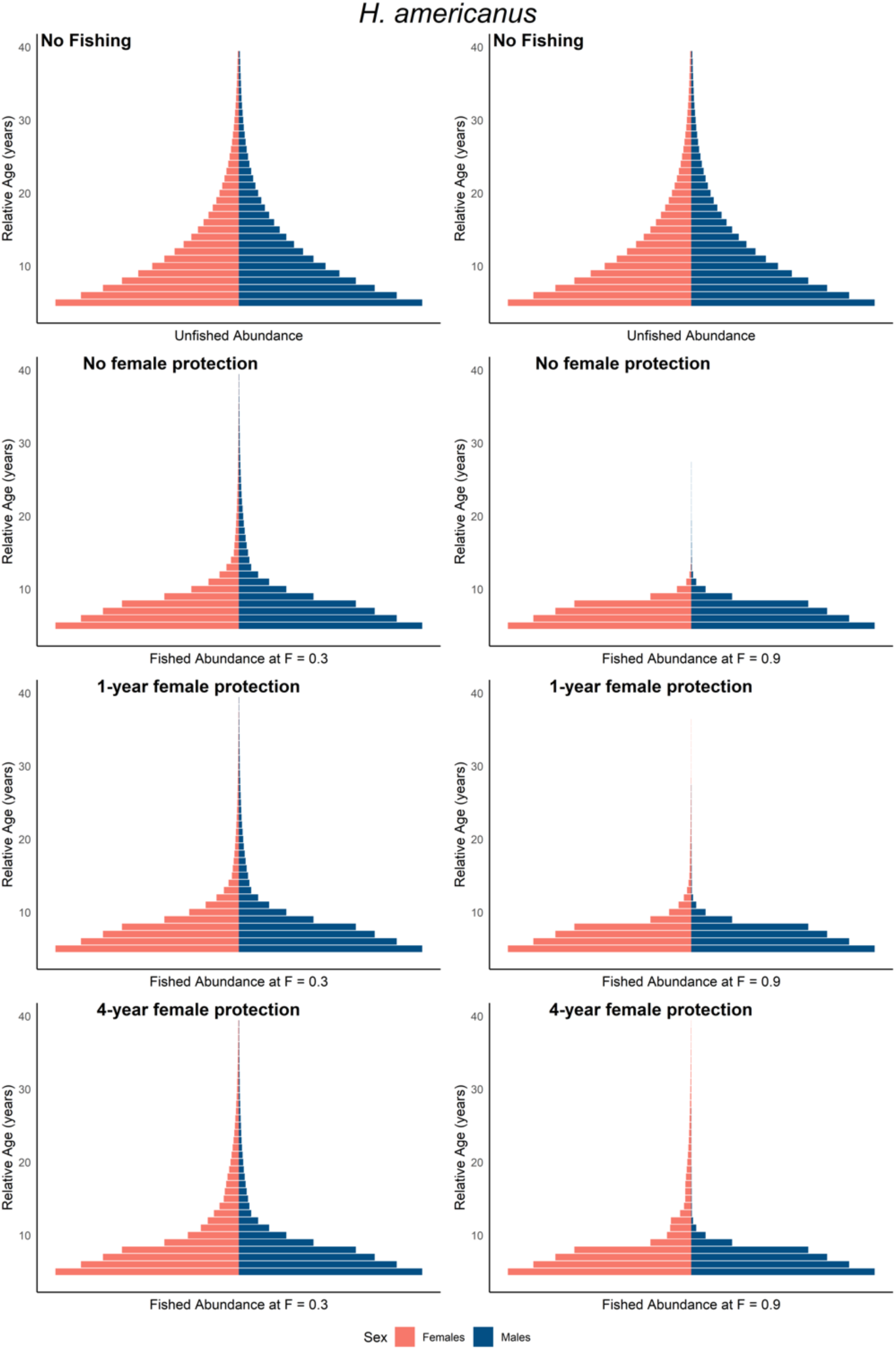
Relative abundance of American lobster individuals at each age in the model, starting at age five. The unfished abundance is shown twice, once in each column for comparison to the different types of female protections under *F* = 0.3 and *F* = 0.9.

**Supplemental Figure 2.**
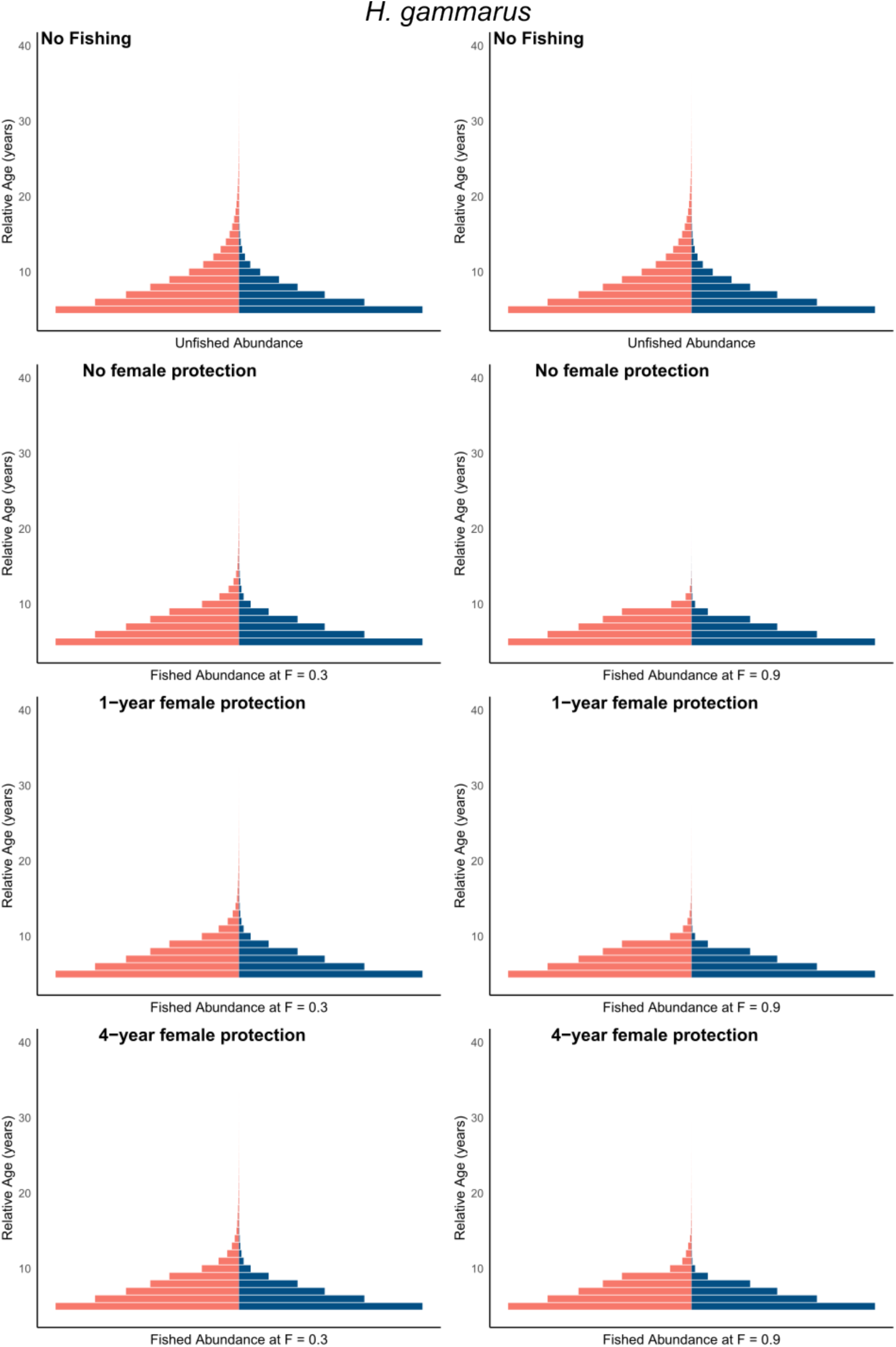
Relative abundance of European lobster individuals at each age in the model, starting at age five. The unfished abundance is shown twice, once in each column for comparison to the different types of female protections under *F* = 0.3 and *F* = 0.9.

